## Supplemental Figures for "Anti-HIV-1 Effect of the Fluoroquinolone Enoxacin and Modulation of Pro-viral hsa-miR-132 Processing"

### Figure S1: CEM-SS Enoxacin effect and qRT-PCR

a)

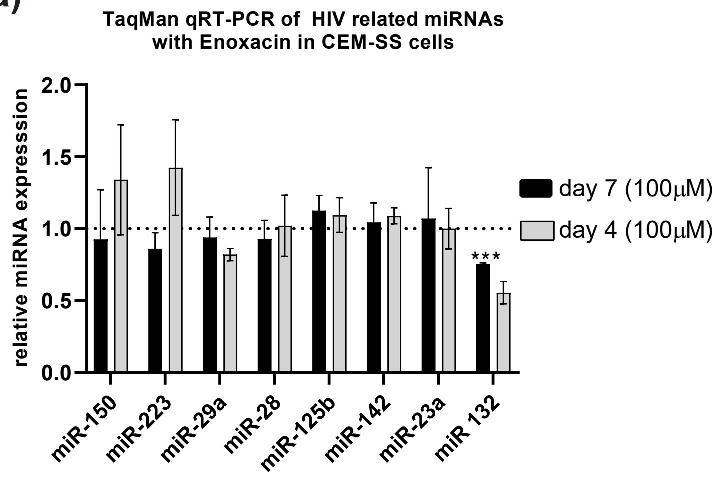


b)

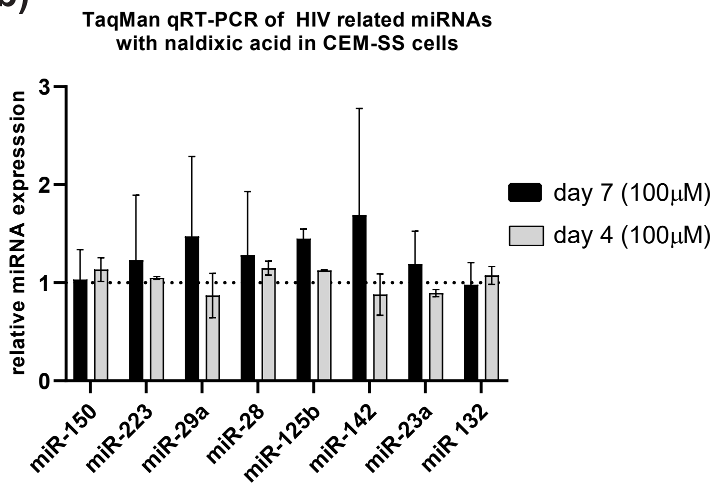


c)

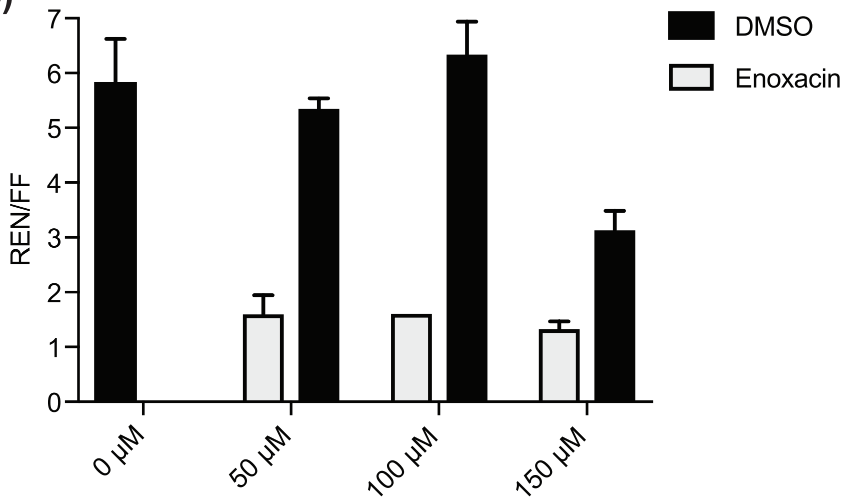

#### Figure S2: Enoaxcin and Nalidixic acid p24 ELISA

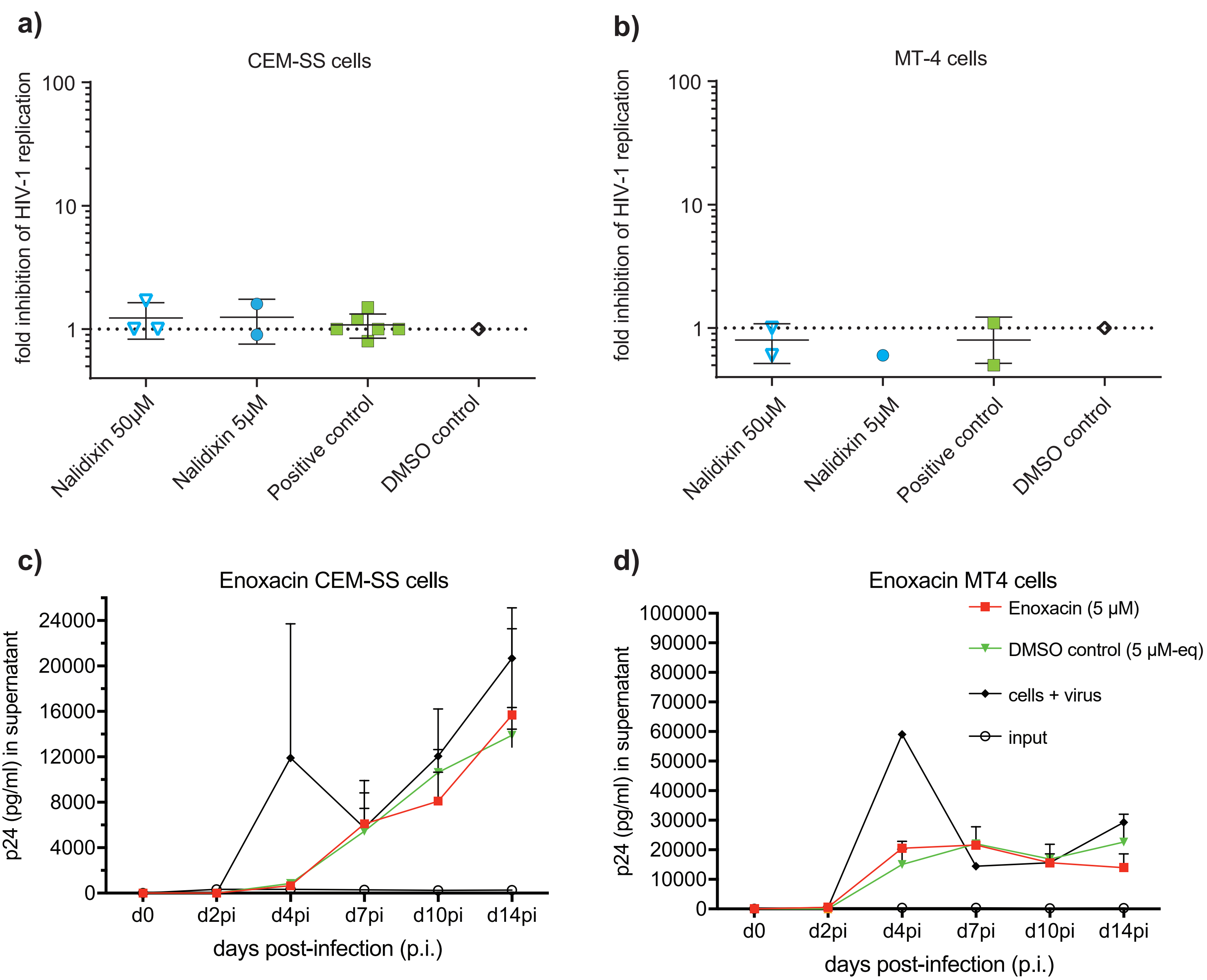

**Figure S3: FACS CEM-SS and MT-4 transfection efficacy**

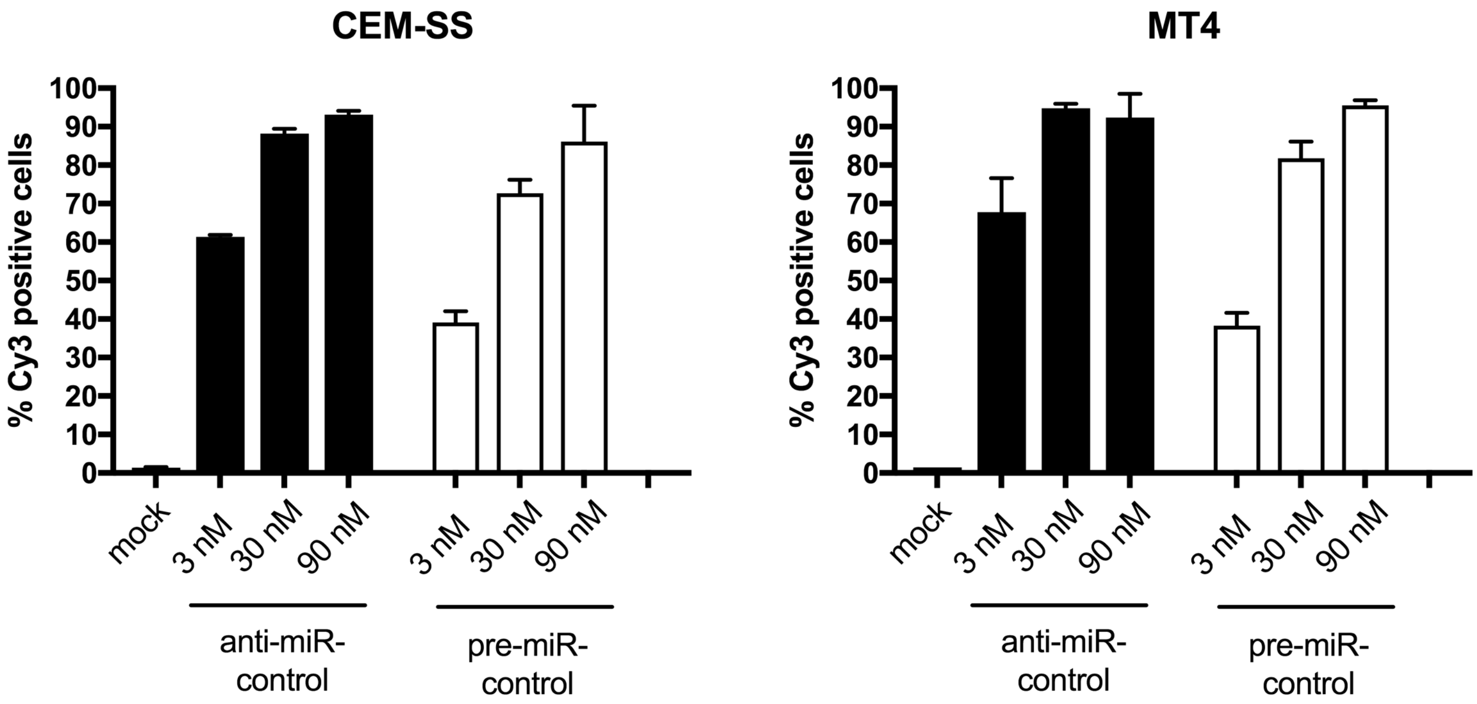

### Figure S4: uncropped 5p Dicer assays and individual quantifications

**a)** miR-132-5p (Rep 1)

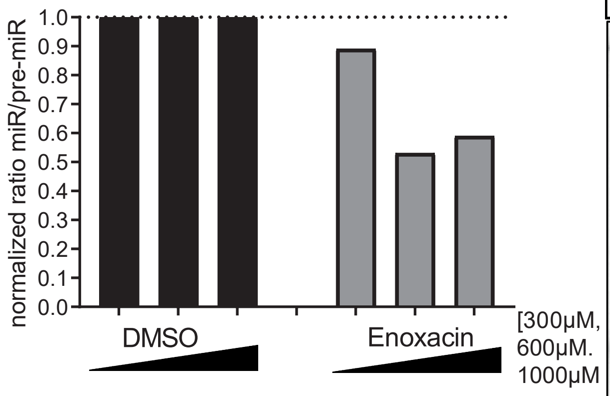


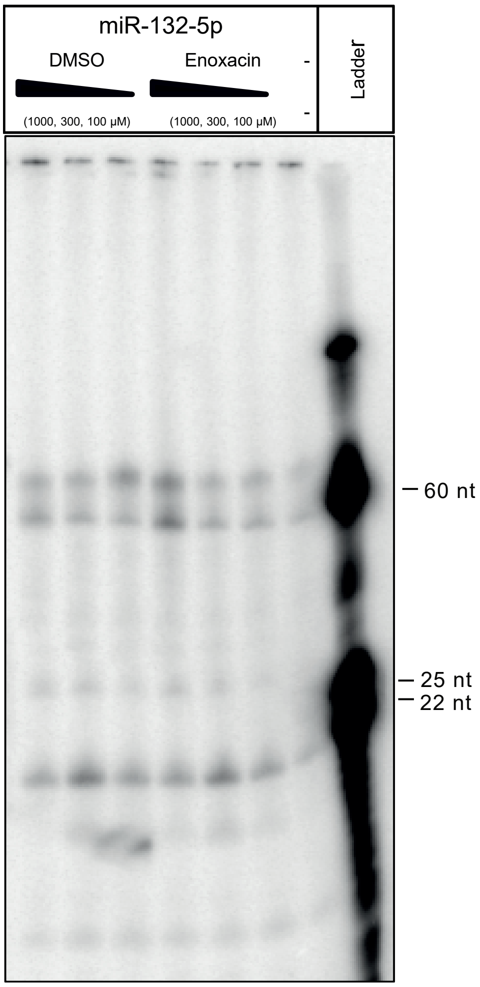


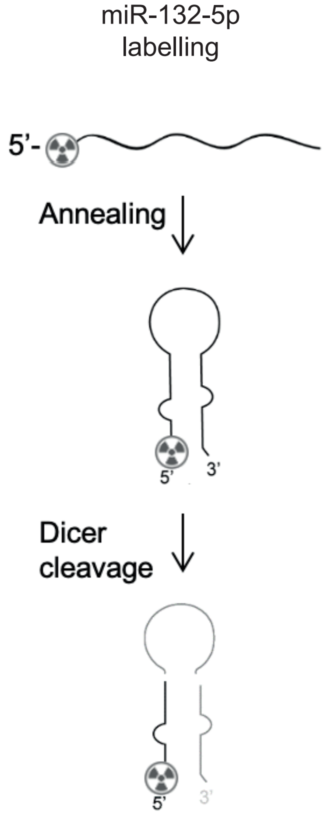


miR-132-5p (Rep 2)

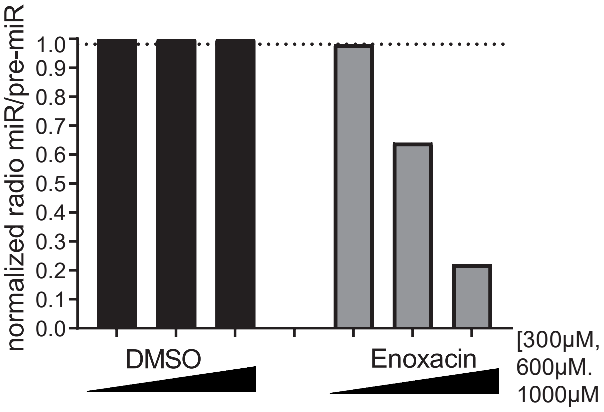


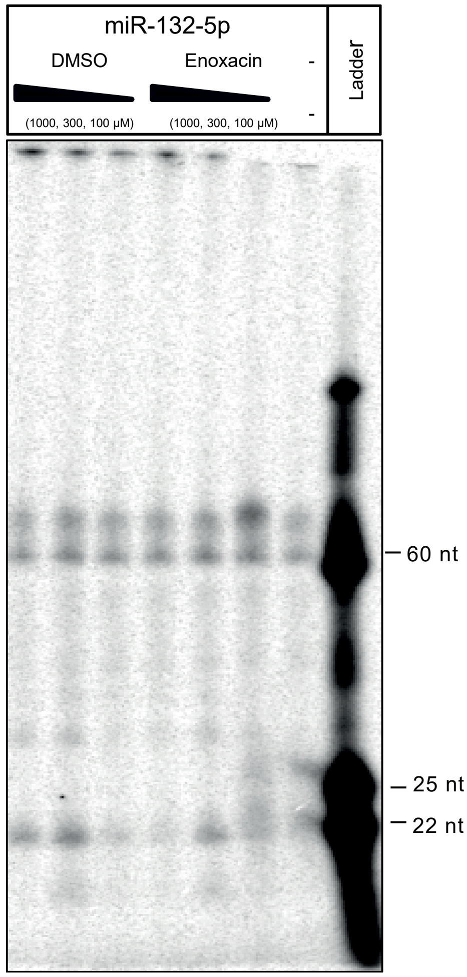

#### Figure S4: uncropped 3p Dicer assays and individual quantifications

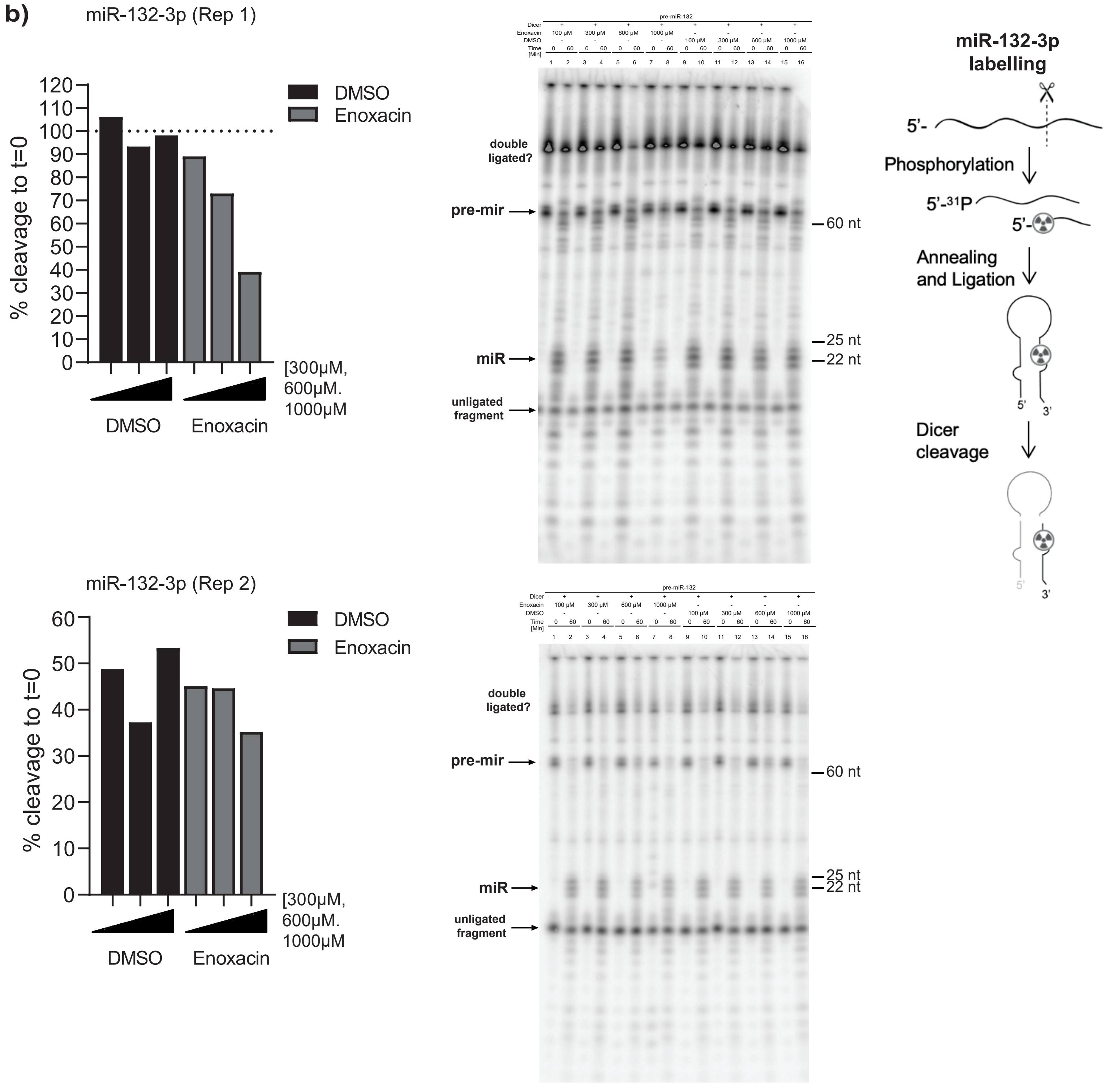

#### Figure S5: SHAPE reactivity plots pre-miR-132

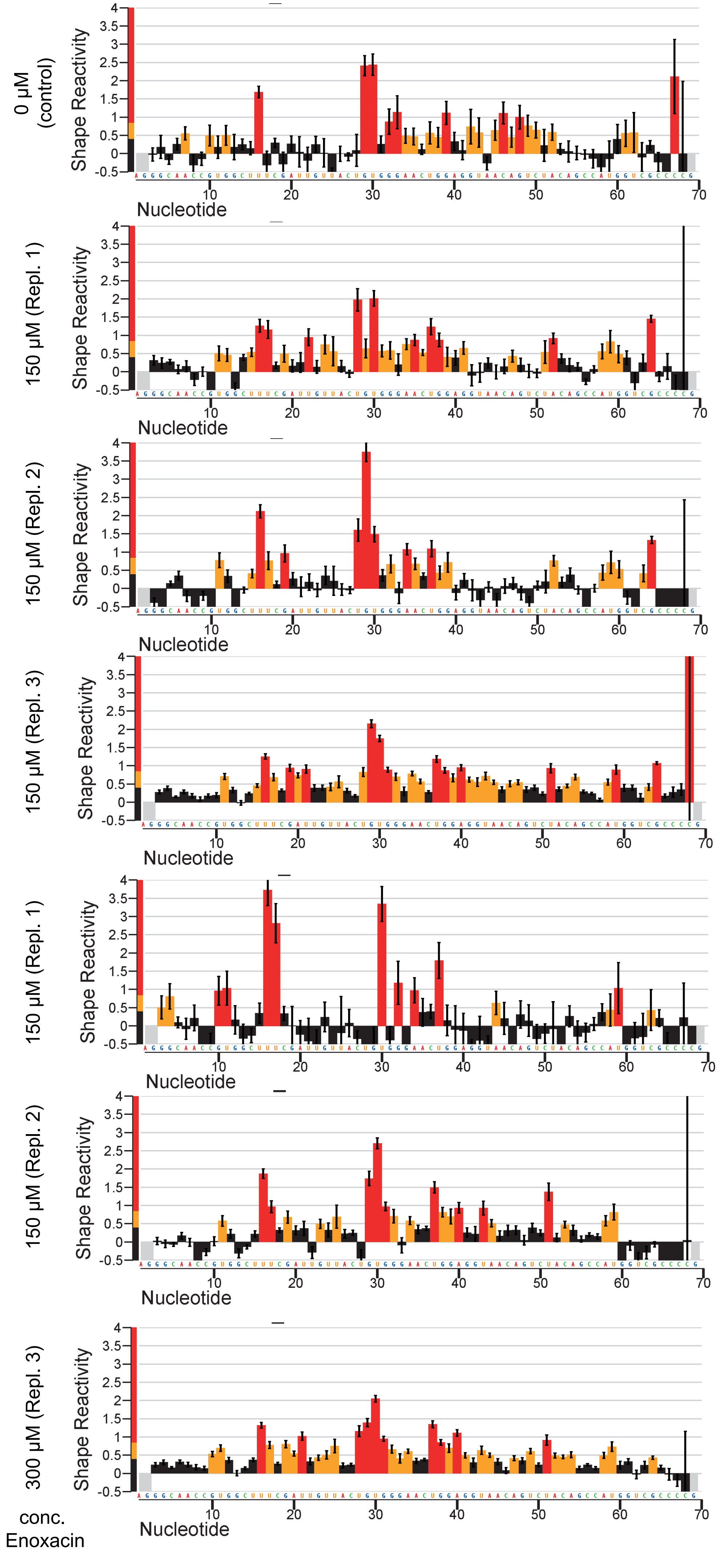
